# Joint Microbial and Metabolomic Network Estimation with the Censored Gaussian Graphical Model

**DOI:** 10.1101/2020.09.07.286880

**Authors:** Jing Ma

## Abstract

Joint analysis of microbiome and metabolomic data represents an imperative objective as the field moves beyond basic microbiome association studies and turns towards mechanistic and translational investigations. We present a censored Gaussian graphical model framework, where the metabolomic data are treated as continuous and the microbiome data as censored at zero, to identify direct interactions (defined as conditional dependence relationships) between microbial species and metabolites. Simulated examples show that our method metaMint performs favorably compared to existing ones. metaMint also provides interpretable microbe-metabolite interactions when applied to a bacterial vaginosis data set. R implementation of metaMint is available on GitHub.

## 1 Introduction

The field of microbiome research is shifting rapidly from cataloging the taxonomic compositions of microbial communities (Huttenhower et al., 2012), to refined technologies that capture strain-level variations or amplicon sequence variants (Lloyd-Price et al., 2017; Callahan et al., 2017; Gilbert et al., 2018), and to multi-omics studies that better capture community functional activity (iHMP Research Network Consortium, 2019). In particular, metabolomics has been extremely useful in explaining microbial functional potential because of its capability in tracking microbially derived metabolites (McHardy et al., 2013; Wu et al., 2016; Jia et al., 2018). Associations between specific microbes and metabolites provide key insights and improved mechanistic models of host-microbe interactions (McMillan et al., 2015; Org et al., 2017; Liu et al., 2017; Lloyd-Price et al., 2019). In practice, the nonparametric Spearman’s rank correlation is often used to quantify the pairwise correlation between microbes and metabolites. However, Spearman’s rank correlation only captures marginal monotonic association, and does not distinguish direct and indirect interactions. In contrast, partial correlations measure conditional dependencies and allow the identification of direct interactions between microbes and metabolites (Gould et al., 2018).

One analytical challenge specific to the microbiome data is the uneven sequencing depths that arise due to differential efficiency of the sequencing process. The total number of reads in a sample is also constrained by the biological specimen at hand and does not reflect the absolute abundance present in the ecosystem. A common practice to address this issue is to transform the raw counts into relative abundances by normalizing over the total sequencing reads in each sample. In other words, raw sequencing counts are transformed into proportions of different microbes whose sum has to be one, also known as compositional data. Several lines of work have been proposed to model marginal and/or conditional microbial interactions from compositional data. For example, SparCC (Friedman and Alm, 2012) and CCLasso (Fang et al., 2015) both estimate the linear Pearson correlations between log transformed counts. A major limitation of marginal association measures such as the Pearson correlation is that they cannot distinguish between direct and indirect relationships (de la Fuente et al., 2004). To address this issue, SPIEC-EASI (Kurtz et al., 2015) learns the conditional dependencies between pairs of microbes while adjusting for effects from other species in the analysis. This is achieved by estimating the inverse covariance of the centered log-ratio (clr) transformed data using e.g. the graphical lasso algorithm (Friedman et al., 2008). Fang et al. (2017) assumes the observed relative abundances follow the logistic normal distribution and proposed a Majorization-Minimization algorithm for learning the conditional dependence relationships among microbes.

Many of the aforementioned methods are specific to microbiome data, and are not directly applicable for joint analysis of microbiome and other omics data types. One naive approach for joint estimation is to apply the graphical lasso algorithm directly to clr transformed microbiome and metabolomic data. However, as illustrated in Figure 1, the Gaussian graphical model may be a poor fit for microbiome data because the marginal distributions of transformed raw counts are in fact highly skewed and often zero inflated.

**Fig. 1.**
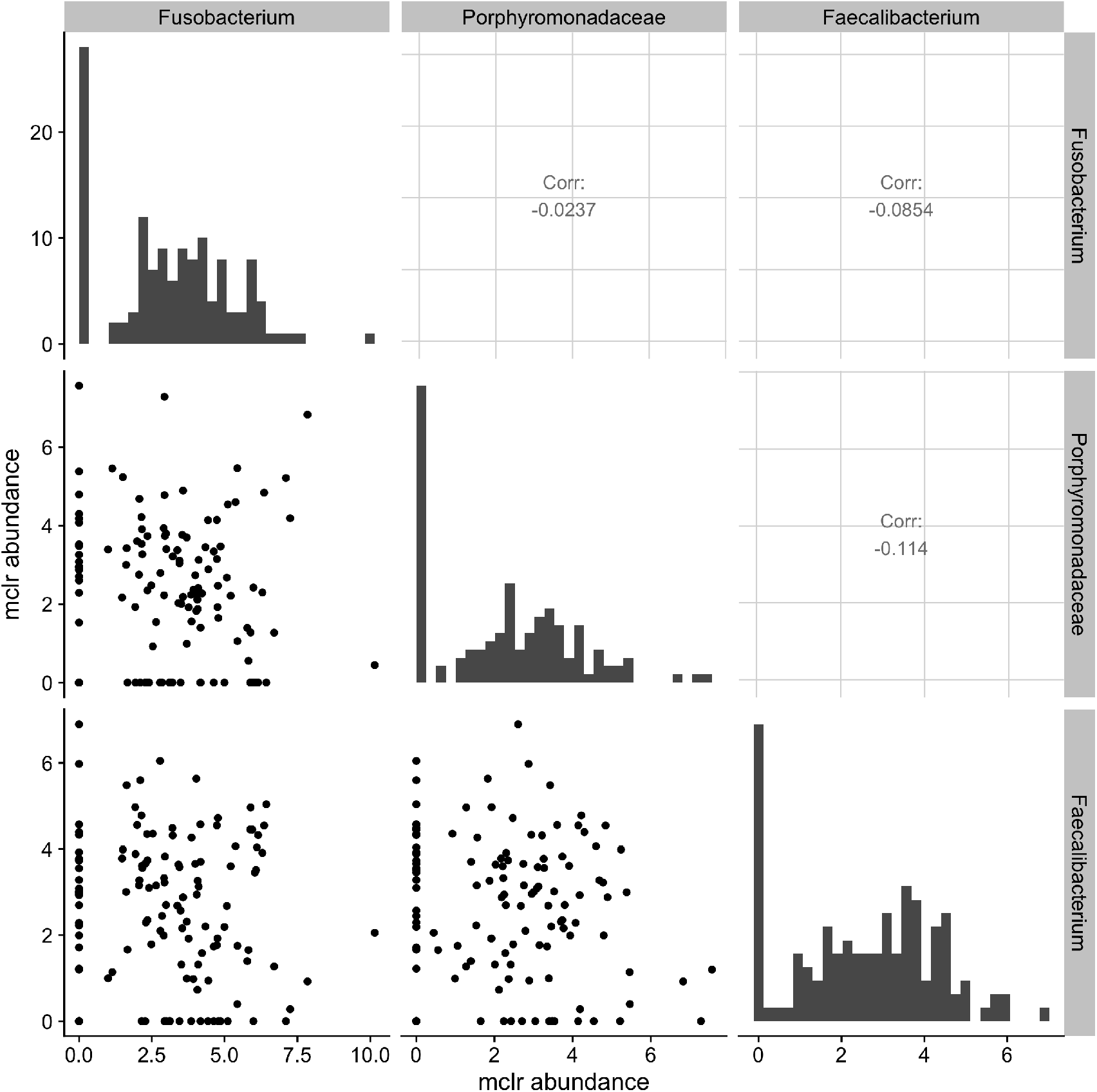
Scatter plots of the modified centered log-ratio (mclr) transformed abundances of 3 bacterial species in the vaginal microbiome data from McMillan et al. (2015). Marginal distribution of each species is illustrated along the diagonal. The upper panels show the Pearson correlations between pairs of species.

This motivates the need for new statistical methodology that can accommodate both microbiome and metabolomic data while accounting for the zero inflation in microbial abundance. Some zero values are sampling zeros that arise due to limited sequencing depths, whereas others are biological zeros that indicate complete absence of a species (Kaul et al., 2017). Silverman et al. (2018) in an unpublished manuscript illustrated that biological zeros in many applications can be approximated as sampling zeros because they both represent a truly low abundance. In this paper, we treat the observed zeros as due to undersampling, and propose a censored Gaussian graphical model (cGGM) to infer the conditional dependencies among microbes and metabolites. Specifically, let ***W*** = (*W*_1_, …, *W*_*q*_)^T^ with *W_j_* > 0 for all *j* be the latent variables, called *the basis*, that represent the true absolute abundance for each species. Due to undersampling and uneven sequencing depths, the observed abundance ***R*** is related to ***W*** via

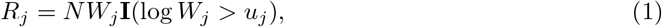

where *N* > 0 is a scaling factor that may depend on ***W***, *u_j_* is a constant which indicates the limit of detection for the *j*-th variable, and **I**(*·*) is the indicator function. The censoring value *u_j_* may be known from the experiment or estimated from data. To adjust for the uneven sequencing depths, we apply the modified-clr (mclr) transformation to ***R***, which transforms all non-zero counts using the usual clr and shifts all transformed values to be strictly positive (Yoon et al., 2019). The diagonal panels in Figure 1 show the histograms of mclr transformed abundances. Compared to the usual clr transformation that requires a pseudo count when dealing with zeros, mclr preserves the ranking of observed counts across multiple samples and is less biased towards rare species (Yoon et al., 2019). Denote ***X***_1_ = mclr_*ε*_(***R***) the resulting vector after mclr transformation with parameter *ε*, which we elaborate in Section 2.3. Let ***X***_2_ = (*X*_*q*+1_, …, *X_p_*)^T^ denote the log transformed concentration measures from *p − q* (*p > q*) metabolites. A natural model for integrating microbiome and metabolomic data is to assume that ***X***_1_ and ***X***_2_ follow a censored multivariate normal distribution with mean ***μ*** and covariance *Σ*. Zero entries in the inverse covariance matrix *Ω* = *Σ*^−1^ capture the conditional independence relationships among the microbes and metabolites.

The problem of inferring the joint microbe-metabolite network thus reduces to estimating *Ω* from *n* independent and identically distributed observations on (***X***_1_, ***X***_2_). We provide metaMint, which is based on estimating each pair of marginal correlations with maximum likelihood. Given the estimated correlation matrix, metaMint uses the graphical lasso to recover the conditional dependencies between microbes and metabolites (direct interactions). We compare our method with several existing approaches in simulations, and show that metaMint outperforms the others in network structural recovery and accuracy of estimating the inverse covariance matrix. When applied to a real data on bacterial vaginosis (McMillan et al., 2015), the integrated network reveals biologically relevant microbe-metabolite interactions and also identifies novel interactions that may serve as potential biomarkers for diagnosis and treatment of bacterial vaginosis.

The censored multivariate normal distribution has been commonly used to analyze environmental data that are often subject to pre-specified detection limits. For example, Hoffman and Johnson (2015), Pesonen et al. (2015) and Jones et al. (2015) studied covariance estimation for left-censored multivariate normal distribution in the classic low-dimensional setting. Recently, Augugliaro et al. (2018) proposed an approximated EM algorithm for inverse covariance estimation in the high-dimensional setting and applied the method to single-cell data. The work by McDavid et al. (2019) was also motivated by single-cell data, but the authors proposed the zero-inflated Gaussian graphical model, which treats zeros as coming from a degenerate point mass at zero instead of being censored. Compared to existing literature, our contribution is a unified model for joint estimation of the integrated microbe and metabolite network in the high-dimensional setting. Our algorithm works well in a variety of scenarios.

The rest of the paper is organized as follows. In Section 2, we describe the censored Gaussian graphical model framework and the proposed algorithm. We present extensive numerical studies in Section 3 and a real data example on bacterial vaginosis in Section 4. We conclude our paper with discussions in Section 5.

## 2 The Censored Gaussian Graphical Model

The censored Gaussian graphical model is suitable for zero-inflated data, which is often the case with microbiome data as shown in Figure 1. In practice, it is reasonable to assume that the observed zeros are due to undersampling or censoring from below.

### Definition 1

A random vector ***X*** is said to follow a censored multivariate normal distribution with mean ***μ*** and covariance *Σ* if there exists constants *u*_1_, …, *u_p_* such that *X_j_* = *Y_j_***I**(*Y_j_ > u_j_*) + *u_j_* **I**(*Y_j_ ≤ u_j_*) where

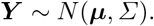

The censoring values ***u*** = (*u*_1_*, …, u_p_*)^T^ are experiment specific and can be inferred from data. For example, one can use the smallest value that occurs more than a pre-specified threshold (e.g. 10%) as an estimate. A pre-specified threshold is necessary to ensure that the smallest value occurs more often than by chance. For zero-inflated microbiome data, the censoring values are set to be 0. When there is no censoring in the *j*-th variable, we set *u_j_* = *−∞*.

The density of the multivariate normal distribution with mean ***μ*** and inverse covariance *Ω* = *Σ*^−1^ is

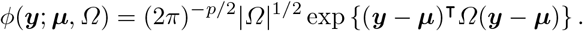

Without loss of generality, let ***X*** = (***X***_*o*_, ***X***_*c*_) where ***X***_*o*_ denotes the uncensored components and ***X***_*c*_ denotes the censored components. Given censoring values ***u*** = (−∞, …, −∞, ***u***_*c*_), the density function of ***X*** is

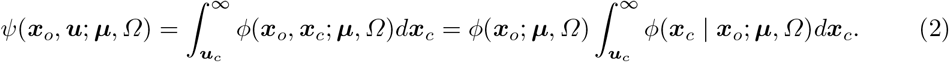

Let {***x***^(1)^, …, ***x***^(*n*)^} denote a set of *n* independent and identically distributed observations on ***X***. In high-dimensional settings, a natural strategy to estimate the inverse covariance matrix is to maximize the *ℓ*_1_ penalized loss function

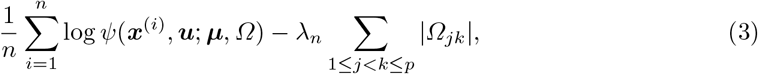

where *λ_n_* is a regularization parameter that controls the sparsity of *Ω*. However, direct optimization of (3) is challenging due to the integral in (2) over a potentially high-dimensional space. Augugliaro et al. (2018) studied a general version of (3) where variables can be left- and right-censored. They proposed to use the EM algorithm to optimize the expectation of the full log-likelihood with respect to the conditional distribution ***X***_*c*_ | ***X***_*o*_. However, exact optimization of the EM algorithm is computationally challenging as it requires the second moment of ***X***_*c*_ | ***X***_*o*_, which is a multivariate truncated Gaussian. The approximation in Augugliaro et al. (2018) is adapted from Guo et al. (2015) and only works well when the inverse covariance matrix is very sparse or the regularization parameter *λ_n_* is large.

### 2.1 A direct estimator via marginal correlations

Our proposal metaMint is based on estimating the marginal correlations directly. A similar idea was used to estimate the correlation matrix of ordinal graphical models (Suggala et al., 2017), where the authors showed that the direct estimator achieves more accurate estimation of the inverse covariance matrix compared to the approximated EM approach in Guo et al. (2015).

The first step in metaMint is to estimate the marginal distribution for each variable, which can be done by fitting a univariate Tobit model (Tobin, 1958) and has been implemented in the R package `censReg` (Henningsen, 2010). Let 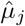 and 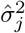 be, respectively, the estimate of the mean and variance for the *j*-th variable. It can be shown that 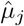 is a consistent estimate of *μ_j_*, and 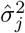 is consistent for 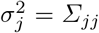. To find the empirical covariance matrix 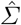, it suffices to estimate each pairwise correlation.

Suppose we have two variables *X_j_* and *X_k_* (*j < k*). If no observation is censored, it is straightforward to estimate their correlation using the Pearson’s correlation coefficient. In the following, we provide details on correlation estimation when at least one variable is censored.

Consider first the case where both variables *X_j_* and *X_k_* are censored from below with *u_j_* and *u_k_*, respectively. For the *i*-th observation, let 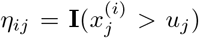 be the indicator function of whether the *j*-th variable is censored. The pairwise joint log-likelihood can be written as a function of the correlation *ρ_jk_*,

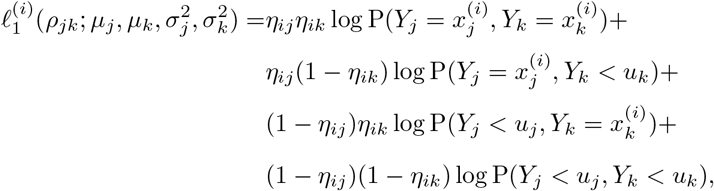

where *Y_j_* and *Y_k_* are bivariate normal with mean (*μ_j_, μ_k_*)^T^ and covariance

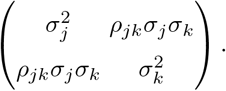

Let *ϕ*(*·*) and *Φ*(*·*) denote, respectively, the density and the cumulative distribution function (c.d.f.) of a standard normal variable. Let the c.d.f. of a bivariate standard normal variable with correlation *ρ* be *Φ*_2_(*u, v, ρ*). The conditional distribution *Y_k_ | Y_j_* = *x*^(*i*)^ is again a normal distribution with mean 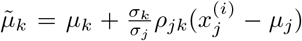 and standard deviation 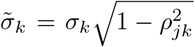. The pairwise joint log-likelihood thus becomes

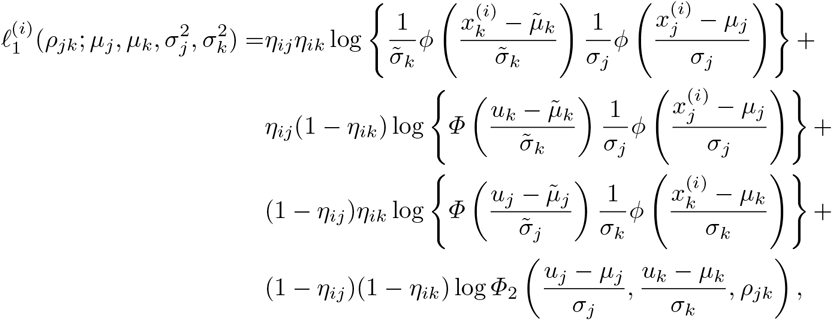

where

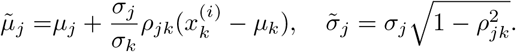

If *u_j_* = *−∞*, this yields a bivariate random vector with only the first variable being censored. Then the joint log-likelihood becomes

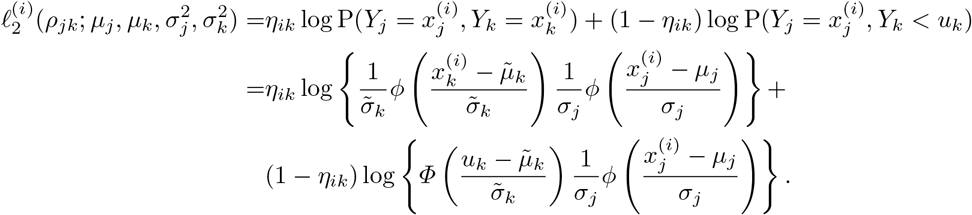

We can solve for *ρ_jk_* as

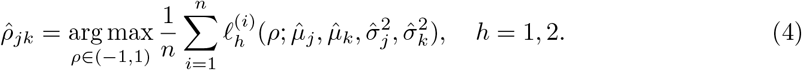

Because entries in 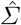 are estimated separately, 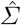 is not guaranteed to be positive semi-definite, which is unsatisfactory because ideally we expect the empirical covariance matrix to be positive semi-definite. One way of bypassing this issue is to use the projection of 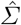 onto a positive semi-definite cone, as done in Fan et al. (2017). In practice, one can calculate the eigen-decomposition of 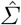 and threshold the negative ones to zero, which yields a new estimator 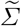.

Given 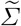, one can apply the graphical lasso algorithm (Friedman et al., 2008)

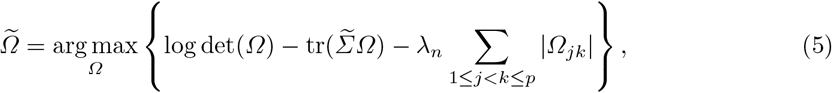

to solve for the inverse covariance matrix *Ω*.

#### Remark 1

The graphical lasso in (5) can be replaced with other algorithms for inverse covariance matrix estimation such as the method by Cai et al. (2011) or its adaptive version (Cai et al., 2016).

metaMint has been implemented in R. In particular, the optimization in (4) is solved using the `optim` function in R, and (5) is solved by the graphical lasso algorithm in the `glasso` package.

### 2.2 Tuning parameter selection

As with other penalization based methods, the proposed algorithm requires the specification of a tuning parameter *λ_n_* that controls the sparsity of the inverse covariance matrix. One can use the cross validation procedure in Guo et al. (2015) or the stability approach in Liu et al. (2010) to select the optimal parameter. In simulations where the ground truth is known, model selection can also be done by maximizing the accuracy in network structural recovery. In Section 4 on real data analysis, we used the stability approach in Liu et al. (2010).

### 2.3 The modified centered log-ratio

The centered log-ratio transformation is often used to transform observed microbial counts to values that are comparable across samples before downstream analysis (van den Boogaart and Tolosana-Delgado, 2013; Gloor et al., 2017; Zhou et al., 2019). Let 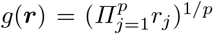 denote the geometric mean of ***r*** = (*r*_1_*, …, r_p_*). The clr of ***r*** is defined as

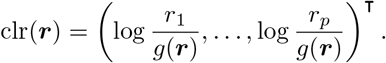

In practice, each sample may consist of many rare species that have zero counts. Thus a pseudo count of 0.5 or 1 is often added to all counts before clr is applied. However, this practice may unfairly bias rare species and impact the accuracy in correlation estimation. The modified centered log-ratio (mclr) (Yoon et al., 2019) attempts to address this limitation by transforming the non-zero counts with the usual clr and shifting all transformed values to be strictly positive.

Without loss of generality, let 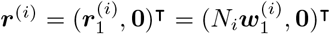 where only components in 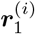 (and 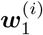) are positive. Although the sample-specific scaling factor *N_i_* does not affect the relative abundances in sample *i*, it captures the variation among total sequencing reads. For example, Vandeputte et al. (2017) observed up to tenfold differences in the total microbial loads after correcting for microbial cell counts. We define mclr_*ε*_(***r***^(*i*)^) as 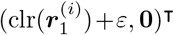, where the constant *ε* is set to be 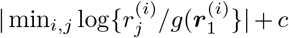 and *c* > 0 is a small constant used to differentiate small positive counts from observed zeros. The resulting mclr_*ε*_(***r***^(*i*)^) is independent of the scaling factor *N_i_*, because 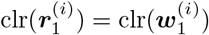. However, adding a pseudo count to zeros and applying clr may introduce unnecessary bias towards zero counts. Figure 2 illustrates the marginal distributions of the genus *Fusobacterium* after the two transformations. Compared to clr, the mclr preserves the relative ranking of all counts while adjusting for the total sequencing depths.

**Fig. 2.**
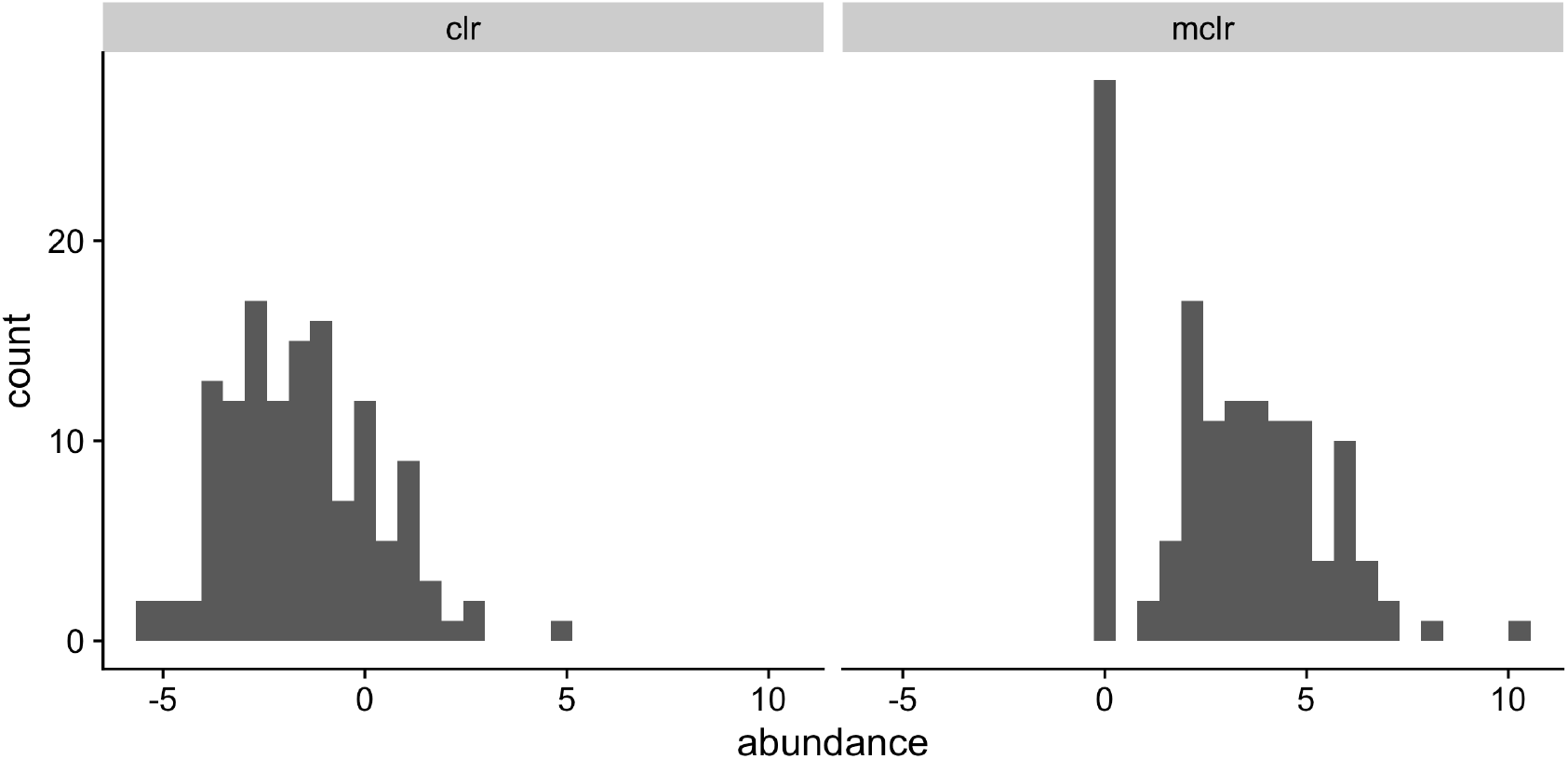
Marginal distributions of the Fusobacterium genus after the clr (left) and mclr (right) transformation to the 131 observations used in McMillan et al. (2015). A pseudo count of 0.5 was added to all counts in order to apply the clr transformation, whereas *c* = 0.1 for mclr.

Lastly, it is worth mentioning that mclr defined above is equivalent to transforming the relative abundances as done in Yoon et al. (2019). To see this, let the relative abundance ***z***^(*i*)^ be defined such that 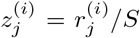, where 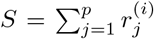. Moreover, we can write 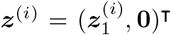 such that only components in 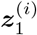 are positive. For any 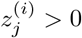,

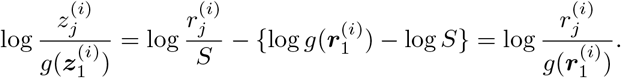

In other words, mclr is scale-invariant.

## 3 Simulation Studies

### 3.1 Model setup

We first generated ***y***^(*i*)^ (*i* = 1, …, *n*) from a multivariate normal distribution with mean ***μ***_0_ and inverse covariance *Ω*_0_. The mean parameter ***μ***_0_ was generated uniformly from [−0.5, 2] to reflect the heterogeneity in abundances of microbial sequences and metabolites. To generate the inverse covariance matrix *Ω*_0_, we considered the following network models, each with *p* nodes:

1. Scale-free network. This network was generated using the Barabasi-Albert algorithm (Barabási and Albert, 1999) and has (*p* − 1) edges. The left panel of Figure 3 illustrates a scale-free network.
2. Erdős-Rényi random graph (Erdős and Rényi, 1960). This network has *p* edges, as illustrated in the middle panel of Figure 3.
3. Nearest-neighbor network. We constructed this network using the same procedure described in Guo et al. (2015), where we uniformly sampled *p* points on a unit square and linked any two points that are 5 nearest neighbors of each other in terms of their Euclidean distances. This network has about 2.5*p* edges. The right panel of Figure 3 illustrates one realization of a sparse network generated with 2 nearest neighbors.

**Fig. 3.**
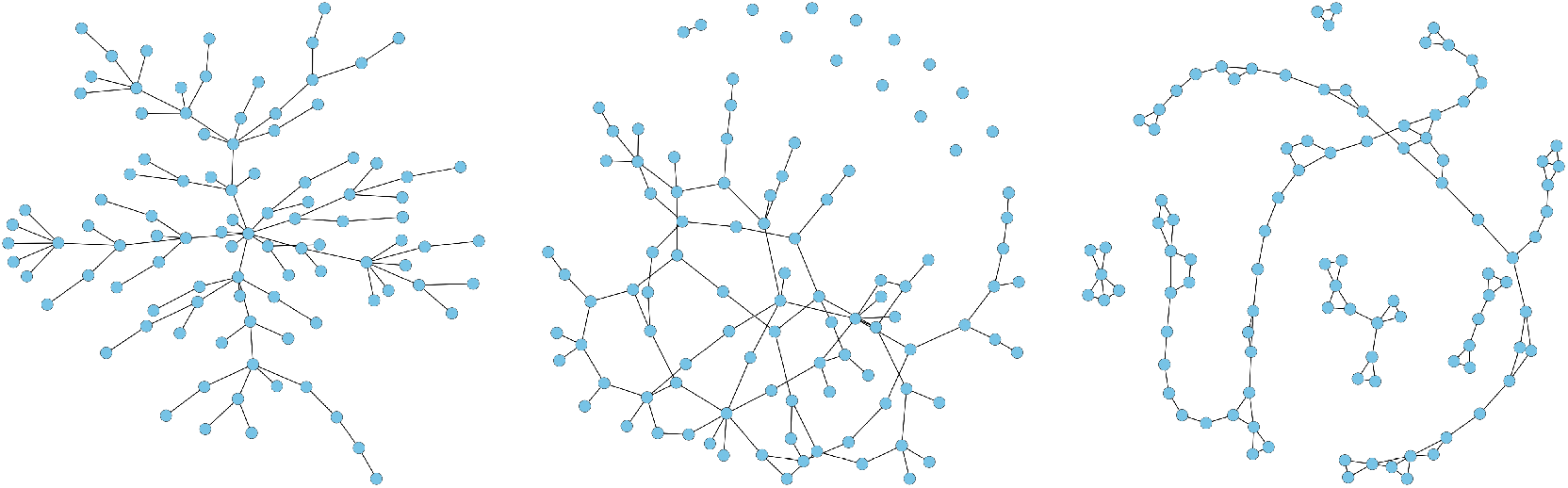
Illustration of graphs used in our simulations (*p* = 100): scale-free graph (left), random graph (middle) and nearest-neighbor graph (right).

Given the network topology, the off-diagonal entries in *Ω*_0_ were generated uniformly from [−1, −0.5] ∪ [0.5, 1], with diagonal entries being 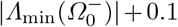. Here 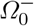 represents the matrix *Ω*_0_ with zeros in the diagonal and *Λ*_min_(*A*) denotes the smallest eigenvalue of *A*. The covariance matrix *Σ*_0_ is then determined by

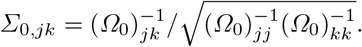

By construction, the diagonal entries of *Σ*_0_ are all 1.

Given the latent ***y***^(*i*)^, the basis vector 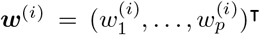 was obtained through the transformation 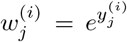. Censored abundances 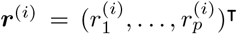 were generated such that

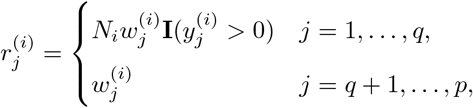

where *N_i_* is generated uniformly between 1 and 10. Here *q* indicates the number of microbes. Only microbiome data are assumed to be censored and compositional in this article, but this assumption can be relaxed in general. In all simulations, we set the constant *c* = 0.1 in the modified clr transformation. Denote 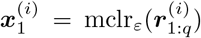 and the observed abundances 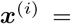 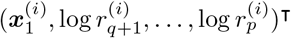.

### 3.2 Results

We compared metaMint with SPIEC-EASI (Kurtz et al., 2015) and gCoda (Fang et al., 2017). The oracle estimator obtained from the latent basis 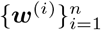 is used as a benchmark, though in practice the oracle is generally unknown. To evaluate the performance of network recovery, we used the receiver operating characteristic (ROC) curve to plot the false positive rate (FPR) against the true positive rate (TPR), defined respectively as,

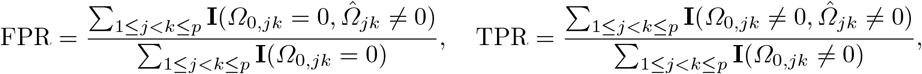

where 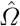 denotes the estimated network. The *F*1 score (van Rijsbergen, 1979), which is between 0 and 1, measures the accuracy of an estimator by summarizing both false positives and false negatives. Larger *F*1 scores indicate better structural recovery. For 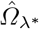 estimated at the optimal penalty parameter *λ*\* selected by maximizing the *F*1 score, we also compared the entropy loss (EL) and Frobenius norm loss (FL) for estimation accuracy:

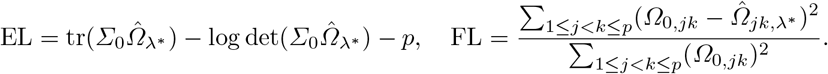

Our first comparison is based on only microbiome data where *p* = *q* = 60 and *n* = 100. In this example, the percentage of zeros per species ranges from 0% to 70%. Input for gCoda is the censored abundance matrix 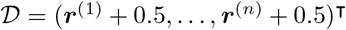. The clr transformation is then applied to each row in 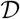 and the resulting matrix is used as input for SPIEC-EASI. Figure 4 shows the ROC curves obtained from different methods across different network models. One can see that SPIEC-EASI and gCoda perform similarly, and both underperform compared to metaMint. Because the nearest-neighbor network is denser, the ROC curves in the right panel of Figure 4 are generally lower compared to their counterparts in other network models.

**Fig. 4.**
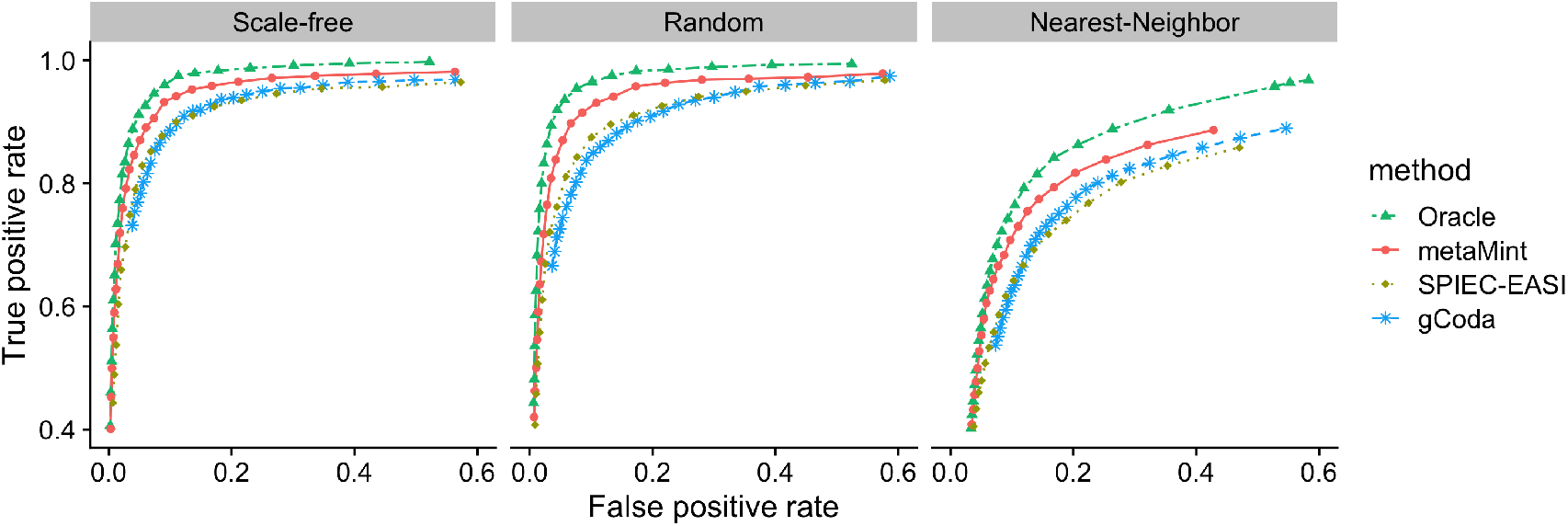
ROC curves for the first study with *p* = *q* = 60 and *n* = 100: Oracle (two-dash line in green), metaMint (solid line in red), SPIEC-EASI (dotted line in brown) and gCoda (dashed line in blue). These results are averaged over 20 replications. metaMint outperforms SPIEC-EASI and gCoda.

In our second study, we look at larger datasets where the number of metabolites is *q* = 100 and the number of microbes is *p − q* = 100. The sample size is *n* = 300. The method gCoda is thus not applicable because it was proposed specifically for microbiome data. Because we only censor microbiome data, the proportion of censored variables in this example is smaller. We first compare different methods in terms of network structural recovery. Figure 5 shows the average *F*1 score of each method across a range of penalty parameters. It can be seen that metaMint has overall higher *F*_1_ scores than SPIEC-EASI, and closely resembles the oracle estimator.

**Fig. 5.**
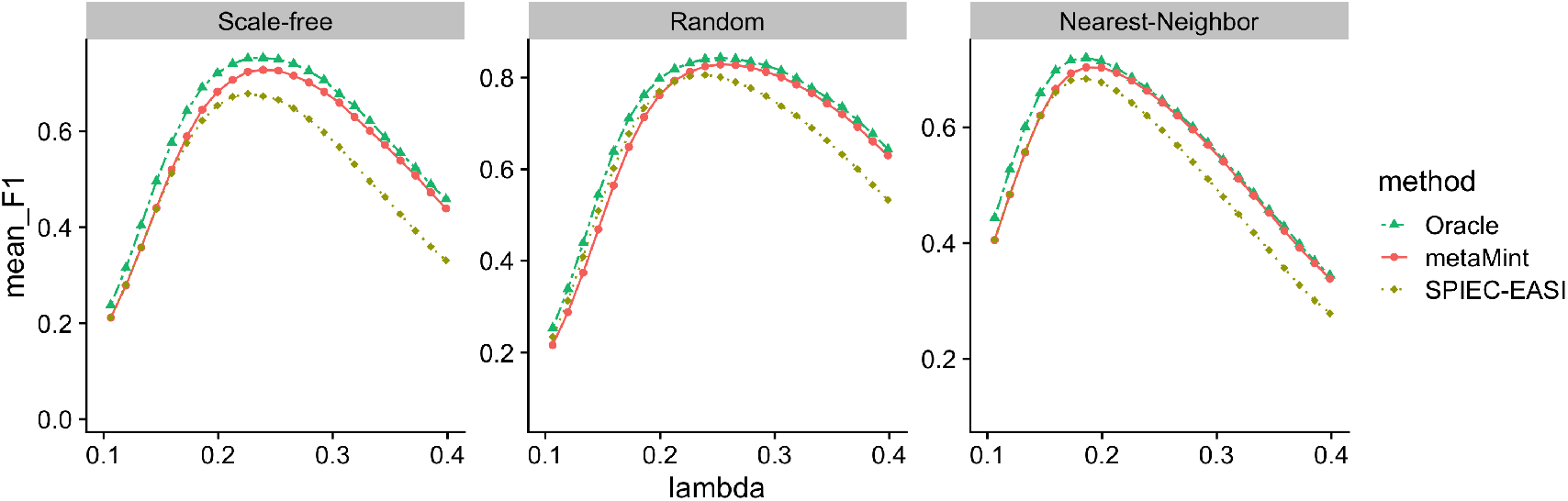
Average *F*1 scores for different methods across different network models over 50 replications in the second simulation study: Oracle (two-dash line in green), metaMint (solid line in red), and SPIEC-EASI (dotted line in brown). gCoda is not applicable in this case because it was specifically for microbiome data. metaMint outperforms SPIEC-EASI.

Since we know the true network structure, we also look at comparisons in terms of inverse covariance estimation accuracy at the optimal penalty parameter selected by maximizing the *F*1 score. As shown in Figure 6, SPIEC-EASI performs the worst in all cases because its entropy and Frobenius norm loss are the largest. It is worth pointing out that there still exists substantial gap in both EL and FL between metaMint and the oracle estimator as a result of censoring. We anticipate that this issue can be partly addressed with increased sequencing depths.

**Fig. 6.**
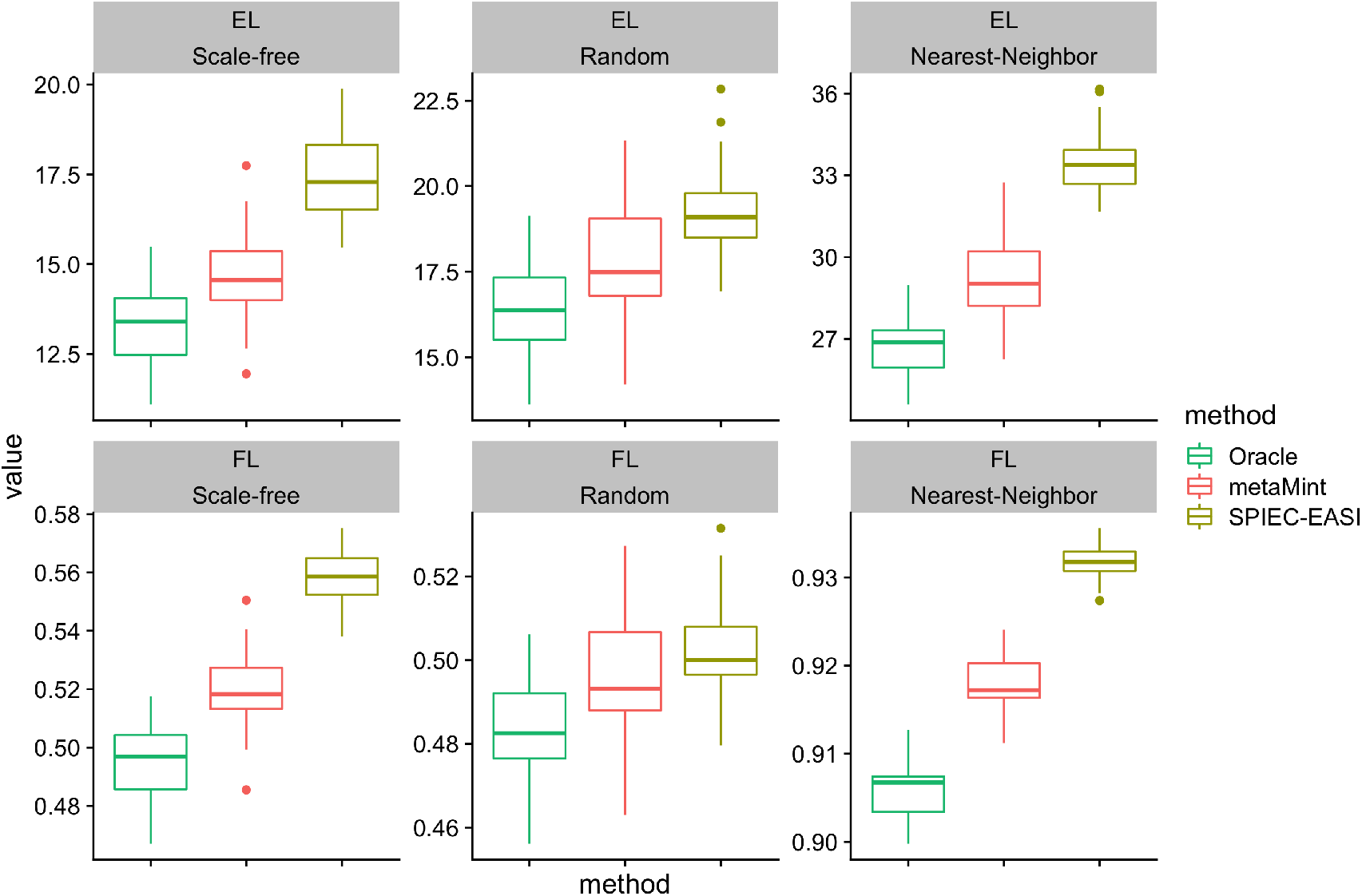
Boxplots showing the entropy loss (EL, top row) and Frobenius norm loss (FL, bottom row) for different methods across different network models over 50 replications in the second simulation study. For each method, the optimal penalty parameter was selected as the one that maximizes *F*1 score. The oracle estimator performs the best, followed by metaMint, and SPIEC-EASI has the largest entropy and Frobenius norm loss.

## 4 Analysis of Bacterial Vaginosis Data

### 4.1 Data description and processing

Bacterial vaginosis (BV) is a common vaginal condition characterized by depletion of specific Lactobacillus species and increased abundance of diverse anaerobic bacteria such as genus *Gardnerella, Prevotella and others* (Fredricks et al., 2005; Ravel et al., 2011). This condition affects an estimated 30% of women at any given time (Koumans et al., 2007), and is associated with increased transmission of HIV and increased risk of preterm labour (Guerra et al., 2006; Atashili et al., 2008). Improved diagnosis and treatment of BV require not only a clearer understanding of the roles of BV associated bacterial species and their interactions, but also a detailed catalog of the interactions between these bacteria and relevant metabolites. We applied the proposed multi-omic approach to a cohort of 131 Rwandan women from McMillan et al. (2015). The microbiome data from sequencing the 16S rRNA gene consist of 51 bacterial species after initial filtering, and the vaginal metabolome determined by GC-MS contains 128 metabolites (see the Methods section in McMillan et al., 2015). One bacterial species is present in only 13 individuals, so we removed this rare species and used 50 taxa in all analysis. Of the 131 women, 79 were normal, 23 were diagnosed with BV, 22 as being intermediate between BV and the normal state, and 7 did not have diagnosis. To account for the different sequencing depths, we applied the clr and modified clr to the microbiome data. Metabolomic data available from McMillan et al. (2015) have already been log transformed. After the mclr transformation, a species is treated as censored at zero if it has at least one zero count. Based on this criterion, 27 of the 50 species are left censored.

We compare metaMint with SPIEC-EASI by applying the former to mclr transformed data and the latter to clr transformed data. At the optimal tuning parameter, which was selected using the stability approach in Liu et al. (2010) with pre-specificed stability threshold *α*, we randomly subsampled 80% of all samples to estimate the network using each method. This procedure was repeated 50 times and an edge selection frequency matrix was constructed such that each entry represents the proportion of times the corresponding edge was present. Only edges with at least 95% selection frequency were kept.

### 4.2 Results

We first compare metaMint and SPIEC-EASI by estimating a single integrated microbe and metabolite network for all subjects at stability threshold *α* = 0.01. Figure 7 presents the joint microbe-metabolite network estimated by the two methods, where the thick black edges are shared between the two methods, blue edges are unique to metaMint, and red edges are unique to SPIEC-EASI. We can see that a majority of edges are shared between the two methods. In particular, both methods reported the conditional association between the genus *Gardnerella* and metabolite *GHB* (6 – 82), and between *Lactobacillus* and *unknown sugar 1* (3 – 165). These two edges are relatively stable and show up in the network for any stability threshold *α ≥* 0.004. Importantly, the interaction between *Gardnerella* and *GHB* was also observed and reproduced experimentally in McMillan et al. (2015). Other notable microbe-metabolite interactions that are unique to each method include *Prevotella* – *unknown sugar 2* (7 – 166) estimated only by metaMint, and *Dialister* – *n-acetyl-putrescine* (10 – 106), *Dialister* – *phenylethylamine* (10 –111) estimated only by SPIEC-EASI. These microbe-metabolite interactions are unique to each method until the stability threshold increases to *α* = 0.02. The differences reported by the two methods are manifestations of the different transformations and whether the model directly accounts for zero-inflation.

**Fig. 7.**
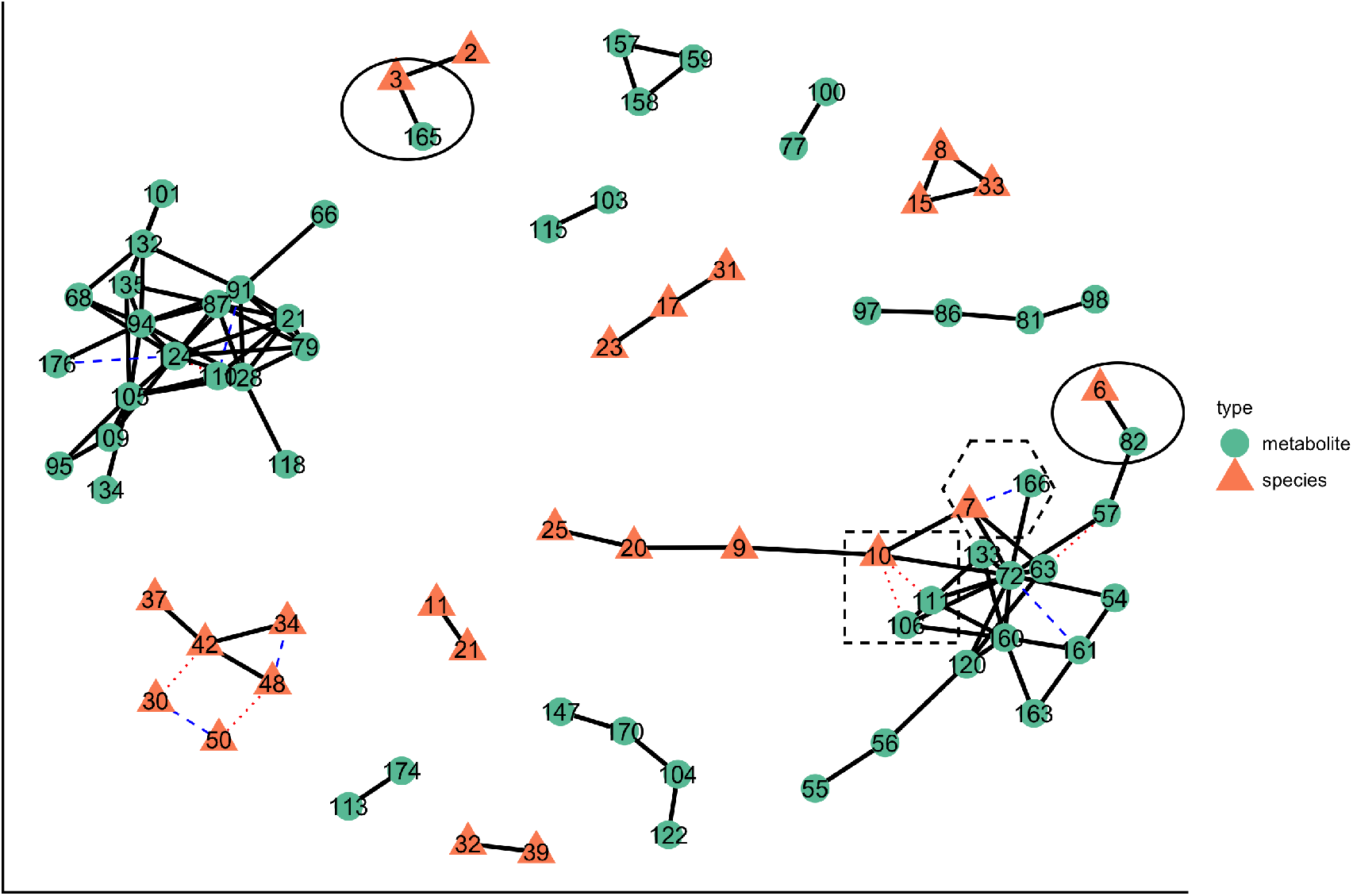
The overlayed microbe-metabolite network in the BV data example estimated from metaMint and SPIEC-EASI. Color and shape of each node indicate whether the node is a metabolite or bacterial species. Thick black edges are shared between the two methods, whereas dashed blue edges are unique to metaMint and dotted red edges are unique to the SPIEC-EASI.

To gain further insights into the roles of these microbe-metabolite interactions, we partitioned all subjects into two groups: the normal group (*n*_1_ = 79) and everyone else (the BV group, *n*_2_ = 52). metaMint and SPIEC-EASI were applied to estimate a network for each group using the same model selection procedure as before. In general, we observe more interactions in the group-specific network estimated by SPIEC-EASI compared to the corresponding network estimated by metaMint. At stability threshold 0.01, no interaction between microbes and metabolites was recovered due to the reduced sample size in each group. As we gradually increase the stability threshold, the first pair of microbe-metabolite interaction unique to the BV group is between *Gardnerella* and *GHB*, and was identified by both metaMint and SPIEC-EASI. Table 1 provides a list of microbe-metabolite interactions that are unique to each group of patients identified by both methods at stability threshold 0.02. It is worth noting that *Gardnerella* – *GHB*, *Prevotella* – *unknown sugar 2*, and *Dialister* – *cadaverine* only show up for the BV group, whereas the interactions between *Lactobacillus* species and several metabolites appear only for the normal group. Abundance of *Lactobacillus* and *Prevotella* has long been used as a diagnostic signature for bacterial vaginosis (Fredricks et al., 2005; Ravel et al., 2011). In addition, McMillan et al. (2015) hypothesized that *Dialister* is responsible for malodor in the vagina. Our analysis may shed light on the mechanistic link between metabolic end products and microbes in vaginal bacterial communities, and provide key guidance regarding the diagnosis and treatment of BV.

**Table 1.** Microbe-metabolite interactions estimated by metaMint and SPIEC-EASI that are unique to each group.

| microbe id | metabolite id | microbe name | metabolite name | Group |
| --- | --- | --- | --- | --- |
| 1 | 62 | Lactobacillus_iners | 2-O-Glycerol-d-galactopyranoside 3 | Normal |
| 2 | 88 | Lactobacillus_crispatus | glyceric_acid | Normal |
| 2 | 125 | Lactobacillus_crispatus | succinate | Normal |
| 3 | 96 | Lactobacillus | malate | Normal |
| 3 | 125 | Lactobacillus | succinate | Normal |
| 6 | 82 | Gardnerella | GHB | BV |
| 7 | 166 | Prevotella | unknown sugar 2 | BV |
| 10 | 72 | Dialister | cadaverine | BV |

## 5 Discussion

The uneven sequencing depths and sparsity in microbiome data present significant challenges in inferring interactions between microbial species and their products. The different sequencing depths imply different levels of uncertainty, but how to handle varying sequencing depths in multivariate statistical analysis remains an unsolved problem (Weiss et al., 2017; McKnight et al., 2019). This paper proposes the censored Gaussian graphical model for joint estimation of microbiome and metabolomic network, which can be used to identify conditional dependencies (direct interactions) between microbial species and metabolites. Key to our proposal is the use of the modified centered log-ratio for transforming the observed microbial counts, which is scale-invariant and preserves the ranking of positive counts relative to zeros. Observed zeros are attributed to undersampling and modeled as due to left censoring. Our method metaMint can be generalized to study other omics data types that fit in the censored Gaussian graphical model framework. Analysis of the bacterial vaginosis data demonstrates that metaMint facilitates the discovery of important microbe-metabolite interactions for diagnosis and treatment of this condition. The data example in Section 4 has about 50% censored variables, although 11 of them have less than 10% zero counts. As we move into high-resolution studies which collect microbiome data at the strain or amplicon sequence variant level, our model that explicitly accounts for observed zeros may exhibit more advantage over existing methods.

From a methodological perspective, metaMint estimates the correlations in a marginal manner, which may not be optimal because marginal approaches ignore the fact that the correlation matrix is positive semi-definite. Augugliaro et al. (2018) proposed an approximated EM algorithm that jointly estimates all entries in the correlation matrix, however their method only works well under specific settings and there is a lack of theoretical understanding about the resulting estimator. Obvious but non-trivial extension is to explore computationally and statistically efficient alternatives that jointly estimate all entries in the correlation matrix.

Our model is related to but substantially different from the zero-inflated Gaussian graphical model in McDavid et al. (2019). While our model assumes the observed zeros are due to undersampling, McDavid et al. (2019) uses a two-part Hurdle model that treats all zeros as structural. The multivariate Hurdle model consists of an Ising model that captures the discrete part and a Gaussian graphical model that describes the continuous part if the hurdle is passed. When the study design favors the two-part process, as is the case in single cell RNA-seq analysis, the multivariate Hurdle model should be considered. On the other hand, the censored Gaussian graphical model is simpler and works well if the study design favors sampling zeros and/or structural zeros can be reasonably approximated as sampling zeros (Silverman et al., 2018).

It is worth pointing out that the observed data defined in (1) are continuous-valued. In this paper, we have made the simplifying assumption that the observed counts can be approximated by a log-normal distribution with left-censoring. An alternative approach is to analyze observed counts directly while still treating zeros as due to left-censoring. In the regression setting, Clark et al. (2017) provided a general framework that uses a latent continuous variable to model observed species abundance, which can be presence/absence, continuous abundance, ordinal counts, or counts that are subject to a total sum constraint. It would be interesting to see if similar ideas can be used to model interactions between microbial species and other molecules.

## Acknowledgements

J. Ma is partially supported by NIH 1R01GM129512-01. The author would like to thank three anonymous referees for their constructive comments and suggestions.

